# Elevating levels of the endocannabinoid 2-arachidonoylglycerol blunts opioid reward but not analgesia

**DOI:** 10.1101/2024.04.02.585967

**Authors:** Arlene Martínez-Rivera, Robert N. Fetcho, Lizzie Birmingham, Jin X Jiu, Ruirong Yang, Careen Foord, Diego Scala-Chávez, Narmin Mekawy, Kristen Pleil, Virginia M. Pickel, Conor Liston, Carlos M. Castorena, Joshua Levitz, Ying-Xian Pan, Lisa A. Briand, Anjali M. Rajadhyaksha, Francis S. Lee

**Affiliations:** Center for Substance Abuse Research, Temple University School of Medicine, Philadelphia, PA, USA; Division of Pediatric Neurology, Department of Pediatrics, Weill Cornell Medicine, New York, NY 10065, USA; Feil Family Brain and Mind Research Institute, Weill Cornell Medicine, New York, NY 10065, USA; Department of Psychiatry, Weill Cornell Medicine, New York, NY 10065, USA; Department of Psychology, Temple University; Neuroscience Program, Temple University, 19122, USA; Department of Anesthesiology, Rutgers New Jersey Medical School, Newark, NJ 07103, USA; Department of Pharmacology, Weill Cornell Medicine, New York, NY 10065, USA; Center for Hypothalamic Research, Department of Internal Medicine, University of Texas Southwestern Medical Center, Dallas, TX 75390, USA; Department of Biochemistry, Weill Cornell Medicine, New York, NY, 10065, USA

## Abstract

Converging findings have established that the endocannabinoid (eCB) system serves as a possible target for the development of new treatments for pain as a complement to opioid-based treatments. Here we show in male and female mice that enhancing levels of the eCB, 2-arachidonoylglycerol (2-AG), through pharmacological inhibition of its catabolic enzyme, monoacylglycerol lipase (MAGL), either systemically or in the ventral tegmental area (VTA) with JZL184, leads to a substantial attenuation of the rewarding effects of opioids in male and female mice using conditioned place preference and self-administration paradigms, without altering their analgesic properties. These effects are driven by CB1 receptors (CB1Rs) within the VTA as VTA CB1R conditional knockout, counteracts JZL184’s effects. Conversely, pharmacologically enhancing the levels of the other eCB, anandamide (AEA), by inhibition of fatty acid amide hydrolase (FAAH) has no effect on opioid reward or analgesia. Using fiber photometry with fluorescent sensors for calcium and dopamine (DA), we find that enhancing 2-AG levels diminishes opioid reward-related nucleus accumbens (NAc) activity and DA neurotransmission. Together these findings reveal that 2-AG counteracts the rewarding properties of opioids and provides a potential adjunctive therapeutic strategy for opioid-related analgesic treatments.

## Introduction

Opioids such as morphine and oxycodone are mainstay analgesics for the management of moderate and severe acute pain. Unfortunately, they are also highly rewarding and can lead to drug dependence, a major contributor to the current opioid epidemic that has led to opioid drug overdoses to become the leading cause of death for Americans under the age of 50 years (1, 2). Thus, the current situation has highlighted the urgent need to develop non-opioid analgesic alternatives. An additional strategy is to develop adjunctive treatments that can specifically attenuate the rewarding, but not the analgesic properties of opioids. One possible avenue is engaging the brain’s endogenous cannabinoid (endocannabinoid; eCB) system that consists of two major neuromodulatory ligands, 2-arachidonoylglycerol (2-AG) and N-arachidonoylethanolamine (anandamide; AEA) (3, 4). These ligands act via the cannabinoid 1 receptor (CB1R) in the brain and regulate the dopaminergic system including the mesolimbic ventral tegmental area (VTA) to nucleus accumbens (NAc) circuit that is at the core of opioid reward (5, 6), with elevations of NAc dopamine associated with the rewarding properties of opioids (7–13).

Endocannabinoid levels are tightly regulated by catabolic enzymes including monoacylglycerol lipase (MAGL) and fatty acid amide hydrolase (FAAH) which respectively hydrolyze 2-AG and AEA (Figure 1A; (14). Recent studies using inhibitors of MAGL and FAAH have revealed an inhibitory role of eCBs on opioid reward. Notably, a dual FAAH-MAGL inhibitor (SA-57), which enhances both 2-AG and AEA levels, reduced heroin self-administration in male mice (15). Similarly, a separate study employing a selective MAGL inhibitor (MJN110), which selectively enhances brain 2-AG levels (16), attenuated morphine place preference in male rats. Presumably these inhibitory effects are via CB1Rs, although this has not been tested. Intriguingly, pharmacological blockade of CB1Rs has a similar inhibitory outcome, leading to the reduction of heroin self-administration (17).

**Figure 1.**
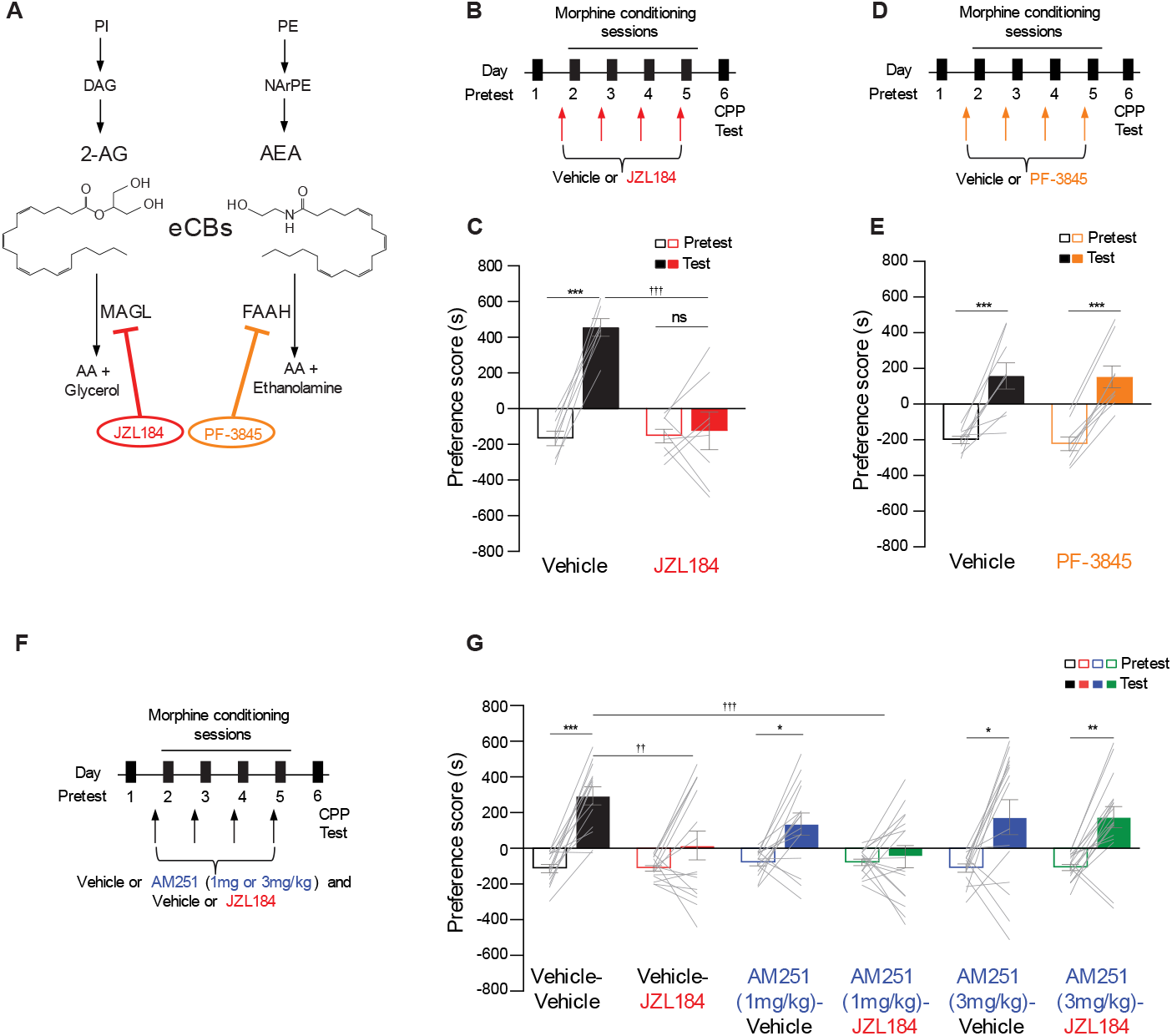
JZL184 attenuates morphine preference via CB1Rs. (**A**) Endocannabinoid signaling pathways regulated by MAGL and FAAH enzymes that are targeted by inhibitors, JZL184 and PF-385, respectively. (**B**) Timeline of behavioral protocol for morphine CPP and JZL184 systemic injections pretreatment. (**C**) Systemic injection of JZL184 prior to each morphine conditioning session abolished morphine CPP (Two-way ANOVA, significant interaction (treatment x day), F_1,26_ =19.77, P=0.0001; Bonferroni post hoc test: Veh: Test vs Pretest P<0.001***, JZL184: Test vs Pretest P>0.9999, Test: Veh vs JZL184 P<0.001†††, Veh n=7, JZL184 n=8). (**D**) Timeline of behavioral protocol for morphine CPP and PF-3845 systemic injections pretreatment. (**E**) Systemic injection of PF-3845 (Two-way ANOVA, main effect of day, F_1,32_=48.49, P<0.0001; Bonferroni post hoc test: Veh: Test vs Pretest P<0.001***, PF-3845: Test vs Pretest P=0.001***, Veh n=9, PF-3845 n=9) had no effect on morphine CPP. (**F**) Timeline of behavioral protocol for morphine CPP and AM251 and/or JZL184 systemic injections pretreatment. (**G**) Systemic injections of the CB1Rs antagonist AM251 (3mg/kg) prior to JZL184 systemic exposure counteracted the JZL184 induced morphine blunted response (Stats were performed by AM251 dose; Three-way ANOVA, significant interaction ((AM251 (3mg/kg) treatment x days x JZL184 treatment), F_1,78_=5.159, P=0.0259*; Bonferroni post hoc test: Veh-Veh: Test vs Pretest P<0.001***, Veh-JZL184: Test vs Pretest P>0.9999, AM251-Veh: Test vs Pretest P=0.0053**, AM251-JZL184: Test vs Pretest P=0.0025**, Test: Veh-Veh vs Veh-JZL184 P=<0.001†††) (AM251 (1mg/kg) significant interaction of AM251 x days, F_1,78_=4.131, P=0.0456*, JZL184 x days, F_1,77_=16.58, P=0.0001***; Bonferroni post hoc test: Veh-Veh: Test vs Pretest P<0.001***, Veh-JZL184: Test vs Pretest P>0.9999, AM251-Veh: Test vs Pretest P=0.0450*, AM251-JZL184: Test vs Pretest P<0.001***, Test: Veh-Veh vs Veh-JZL184 P=<0.001†††), Veh-Veh n=24, Veh-JZL184 n=26, AM251 (3mg/kg)-Veh n=15, AM251 (3mg/kg)-JZL184 n=17, AM251 (1mg/kg)-Veh n=15, AM251 (1mg/kg)-JZL184 n=16). Error bars ± SEM.

The complexity of the eCB system is further emphasized from studies on the effect of AEA and 2-AG on motivated behaviors and the dopamine system. While increasing AEA levels decreased cue-induced reward seeking and NAc dopamine (18), 2-AG has the opposite effect. Enhancing 2-AG with MAGL inhibition (JZL184) enhanced intracranial self-stimulation and food reward seeking as well as NAc dopamine in a CB1R-dependent manner (18).

In light of the divergent results on the role of eCBs on motivated reward related behaviors and opioid reward, our study aimed to investigate the specific impact of enhancing 2-AG and AEA on opioid reward. We additionally assessed the dependency on CB1Rs and examined the effects on neural activity in the NAc and dopamine dynamics. We found that pharmacologically inhibiting MAGL led to a substantial attenuation of the rewarding effects of morphine and oxycodone in male and female mice. Conversely, pharmacologically enhancing the levels of AEA had no effect on morphine reward. In addition, we found that inhibiting MAGL locally in the VTA was sufficient to blunt morphine reward and diminished reward associated NAc activity and DA transmission. In sum, we have identified a unique interaction between the eCB and opioid systems in which 2-AG selectively counteracts the rewarding properties of opioids via disruption of NAc reward-related neural activity and DA transmission while preserving their analgesic effects.

## Results

### Pharmacologically inhibiting MAGL and not FAAH attenuates morphine preference

To study the effect of enhancing eCB levels on the rewarding properties of opioids, we utilized the pharmacological MAGL inhibitor JZL184, which increases 2-AG levels by inhibiting its hydrolysis (19), or the FAAH inhibitor PF-385, which increases AEA levels by inhibiting its hydrolysis (20) (**Figure 1A**). We initially tested the rewarding properties of morphine, a widely-used mu opioid receptor (MOR) agonist with strong reinforcing properties both in humans (21) and animals (22) utilizing the conditioned place preference (CPP) assay, a well-established rodent behavioral proxy for drug reward (23). After an initial CPP pretest on day 1, adult male mice were systemically administered JZL184 (10 mg/kg, intraperitoneal (i.p.); **Figure 1B**) or PF-3845 (10 mg/kg, i.p.; **Figure 1D**) 2 hours prior to each of the four morphine conditioning sessions (days 2-5). Twenty-four hours later mice were tested for morphine preference in the absence of both the inhibitor and morphine (**Figure 1B, 1D**). We found that JZL184 during the conditioning phase of CPP attenuated morphine preference (**Figure 1C)**. A similar effect was observed in females (**Supplementary Fig 1A-B).** In contrast enhancing AEA with PF-3845 at 10mg/kg, i.p. (**Figure 1E**), and 20 mg/kg, i.p. (**Supplementary Figure 1C-D**), had no effect on morphine CPP suggesting that the decrease in morphine preference is unique to 2-AG over AEA. The effect of JZL184 on drug preference was independent of any changes in locomotor activity (**Supplementary Figure 1E**), anxiety-like behavior (**Supplementary Figure 1F**) or depressive-like behavior (**Supplementary Figure 1G**) compared to vehicle treated animals. Additionally, JZL184 on its own had no effect on morphine preference (**Supplementary Figure 1H-I**), and pretreating mice with JZL184 prior to the morphine CPP expression test had no effect either (**Supplementary Figure 1J-K**). The doses and pretreatment time points of JZL184 and PF-3845, both irreversible covalent inhibitors (24, 25) were based on previously published work (26, 27).

Next, as 2-AG acts primarily through CB1Rs (28, 29), we tested if the effect of JZL184 was via CB1Rs. Male mice were pretreated with the CB1R antagonist AM251 (1mg/kg, i.p. or 3mg/kg. i.p. (**Figure 1F-G**), prior to treatment with vehicle or JZL184 during the morphine conditioning phase of CPP (**Figure 1F**). As expected, JZL184 on its own blunted morphine preference (**Figure 1G**; Vehicle-JZL184 vs. Vehicle-Vehicle). Pretreatment with 3mg/kg, i.p. AM251 (**Figure 1G**; AM251 (3mg/kg)-JZL184 vs. Vehicle-JZL184) but not 1mg/kg, i.p. AM251 (**Figure 1G**; AM251(1mg/kg)-JZL184 vs. Vehicle-JZL184) reversed JZL184’s effect on attenuating morphine preference, demonstrating that JZL184 acts via CB1Rs. Of note, AM251 (1 and 3 mg/kg, i.p.) on its own lowered morphine preference, however mice acquired a preference that was not significantly different than vehicle treated mice (**Figure 1G**; AM251-Vehicle vs. Vehicle-Vehicle). Taken together the above findings indicate that JZL184 blunts morphine preference via CB1Rs.

### Pharmacologically inhibiting MAGL attenuates oxycodone preference and self-administration

To test if the effect of JZL184 is generalizable across opioids, we tested its effect on oxycodone CPP and self-administration. First, using CPP (**Figure 2A**), we found that, similar to morphine, repeated dosing of JZL184 during oxycodone conditioning attenuated preference in adult male mice (**Figure 2B**) without altering locomotor activity (**Supplementary Figure 2A**). We next tested the effect of JZL184 on oxycodone self-administration (**Figure 2C**). We chose to use oxycodone for the self-administration studies due to pharmacokinetic differences between morphine and oxycodone in mice. The shorter time to peak and half-life of oxycodone seems to lead to higher consumption and less intoxication in the intravenous self-administration model. Mice were first food-trained on the operant chamber for two days (**Supplementary Fig 2B**). Then, mice were randomly separated into two groups, one pretreated with JZL184 (10mg/kg) and the other pretreated with vehicle before each oxycodone-self administration session (0.25mg/kg/infusion, **Figure 2C**). Animals pretreated with JZL184 self-administered less oxycodone when compared with the control group (**Figure 2D-E**). The percent of correct choices did not differ between groups (**Supplementary Figure 2C-D**). Similar results were obtained with female mice (**Supplementary Figure 2E-J**). These results demonstrate that JZL184 reduces the rewarding and reinforcing properties of oxycodone and that JZL184’s effects might be generalized across different opioids.

**Figure 2.**
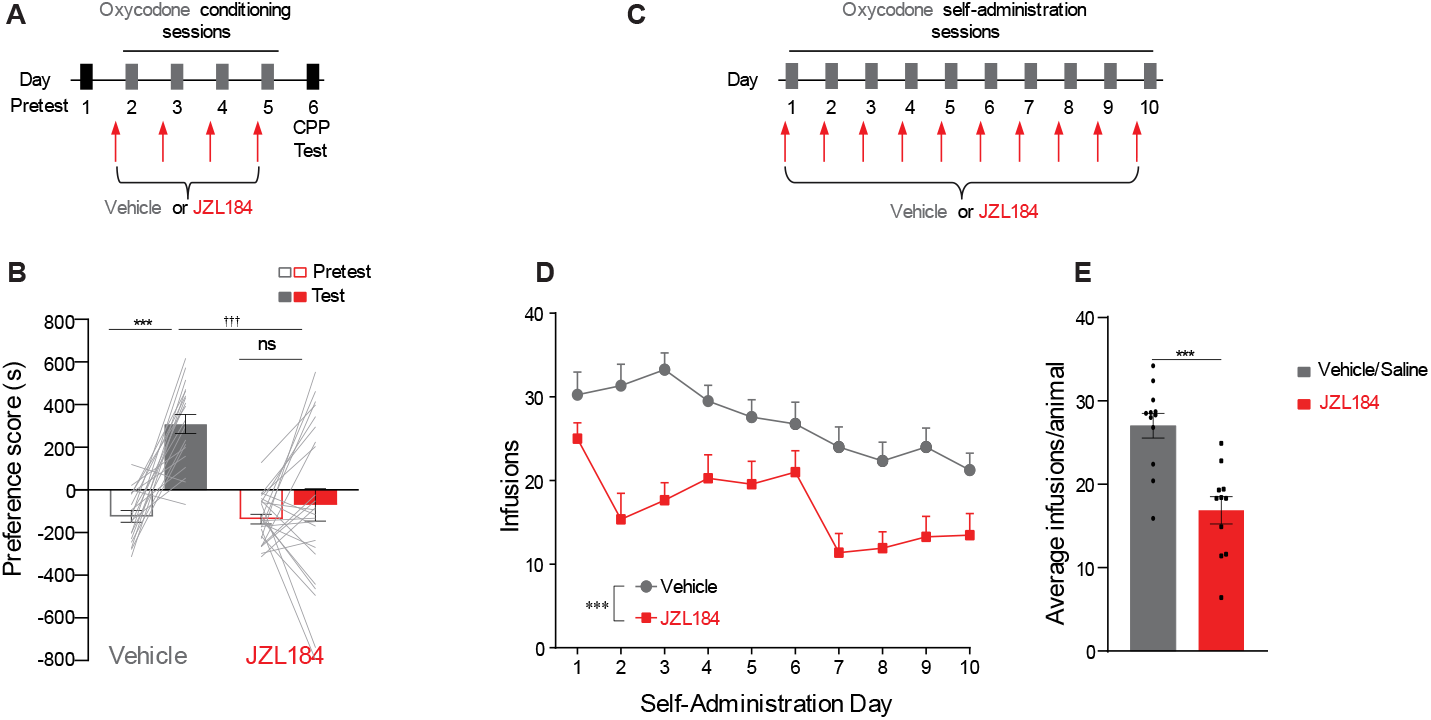
JZL184 attenuates oxycodone preference and self-administration. (**A**) Timeline of behavioral protocol for oxycodone CPP and JZL184 systemic injections pretreatment. (**B**) Systemic JZL184 prior to each oxycodone conditioning session attenuated oxycodone CPP (Two-way ANOVA, significant interaction (treatment x day), F_1,82_ =13.31, P=0.0005; Bonferroni post hoc test: Veh: Test vs Pretest P<0.001***, JZL184: Test vs Pretest no significant, Test: Veh vs JZL184 P<0.001†††, Veh n=19, JZL184 n=24). (**C**) Timeline of behavioral protocol for oxycodone self-administration and JZL184 systemic injections pretreatment. (**D-E**). Systemic JZL184 exposure prior to oxycodone self-administration sessions, attenuated the intake of oxycodone (**D;** Two Way-RM ANOVA, main effect of JZL184 treatment, F_1,21_ =21.43, P=0.0001***, main effect of days, F_2.962,62.20_ =8.236, P=0.0001***, interaction F_9,189_ =1.900, P=0.0541, **E;** Average of total infusions/animal (Ttest, t(_21_) = 4.63, P< 0.0001***, Veh/Saline n=12, JZL184 n=11). Error bars ± SEM.

### Pharmacologically inhibiting MAGL does not alter opioid analgesia

Given the positive effects of JZL184 on attenuating opioid reward, we next queried whether JZL184 would alter the analgesic effects of morphine using the hot plate assay (30) and the radiant-heat tail-flick test, a behavioral assay commonly used to test pain response in animals (31). For the hot plate test, mice were treated with a single dose of JZL184 (10 mg/kg, i.p.) and 2 hours later a morphine dose-response (0.3, 1.0, 3.0 and 10 mg/kg, i.p) test was performed (**Figure 3A**). JZL184 had no effect on morphine analgesia (**Figure 3B top**). The ED_50_ value of vehicle-treated control animals was comparable to that in animals that received a single dose of JZL184 (**Figure 3B bottom panel**). Similarly, in the tail flick test a single dose of JZL184 had no effect on the ED_50_ value of morphine (**Figure 3C-D**) or oxycodone (**Supplementary Figure 3A-B**).

**Figure 3.**
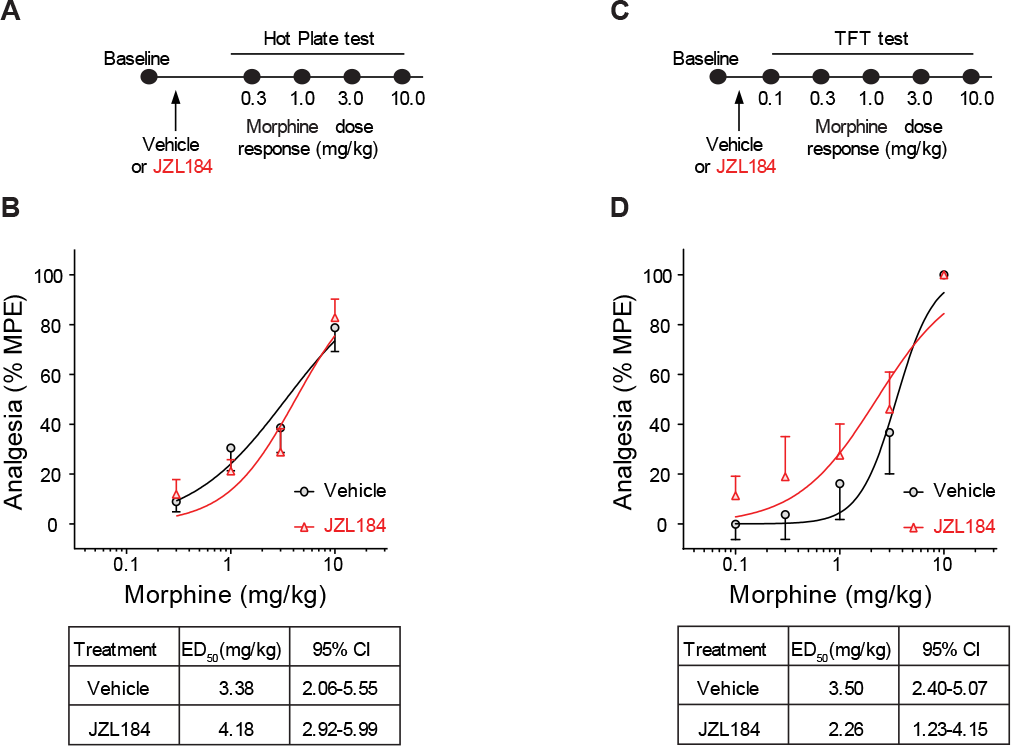
JZL184 has no effect on morphine analgesia. (**A, C**) Experimental timeline of acute systemic vehicle or JZL184 pretreatment prior to the morphine dose response in the hot plate test (**A**), or the tail-flick test (**C**). (**B, D**) JZL184 pretreatment has no effect on morphine-induced analgesia during the hot plate test (Two-way ANOVA, main effect of dose, F_3,68_= 30.65, P<0.001***, Veh n=9, JZL184 n=10), (**D**) or the tail flick test (Two-way ANOVA, main effect of dose, F_4,60_ =22.17, P<0.001***, Veh n=7, JZL184 n=7). Error bars ± SEM.

We additionally examined whether repeated JZL184 administration similar to the treatment regimen used in morphine CPP (**Supplementary Figure 3C-D**) would affect morphine analgesia. For this experiment, male mice were randomly separated into two groups and first subjected to a morphine dose response to measure morphine analgesia (Supplementary **Figure 3D** **middle/bottom**; Pre-Vehicle and Pre-JZL184 groups). Mice then received injections of vehicle or JZL184 (10 mg/kg, i.p.) once a day for four days. Twenty-four hours later mice were re-tested for morphine analgesia (**Supplementary Figure 3D**; Post-Vehicle and Post-JZL184 groups). Similar to the above results, we did not observe significant differences in the ED_50_ values between repeated JZL184 and vehicle pretreated mice. Additionally, enhancing AEA with PF-3845 (10 mg/kg, i.p.; **Supplementary Figure 3E-F**) had no effect on morphine analgesia. Together these data indicate that systemic elevation of 2-AG levels does not impair the ability of opioids to mediate analgesia in mice.

### Inhibiting MAGL in the ventral tegmental area (VTA) blunts morphine preference but not analgesia

Having observed specific effects of inhibiting MAGL on opioid reward but not analgesia, we asked which brain region is involved in this effect. As the VTA is a central site for driving opioid reward (5), we tested whether the systemic effect of enhancing 2-AG was driven by its action locally in the VTA. JZL184 (5 µg/µL per side) was infused bilaterally into the VTA of male mice prior to each of the four morphine conditioning sessions (**Figure 4A-B, Supplementary Figure 4A**) and twenty-four hours later mice were tested for morphine preference in the absence of both JZL184 and morphine. As was seen with systemic delivery (**Figure 1B**), repeated application of intra-VTA JZL184 attenuated morphine preference (**Figure 4C**). Furthermore, consistent with systemic JZL184 treatment, intra-VTA JZL184 infusion had no effect on morphine analgesia (**Supplementary Figure 4B**), locomotor activity (**Supplementary Figure 4B**), anxiety-like (**Figure Supplementary 4C**) or depressive-like behaviors (**Supplementary Figure 4D**), demonstrating that 2-AG is acting via effects on the VTA to specifically blunt opioid reward.

**Figure 4.**
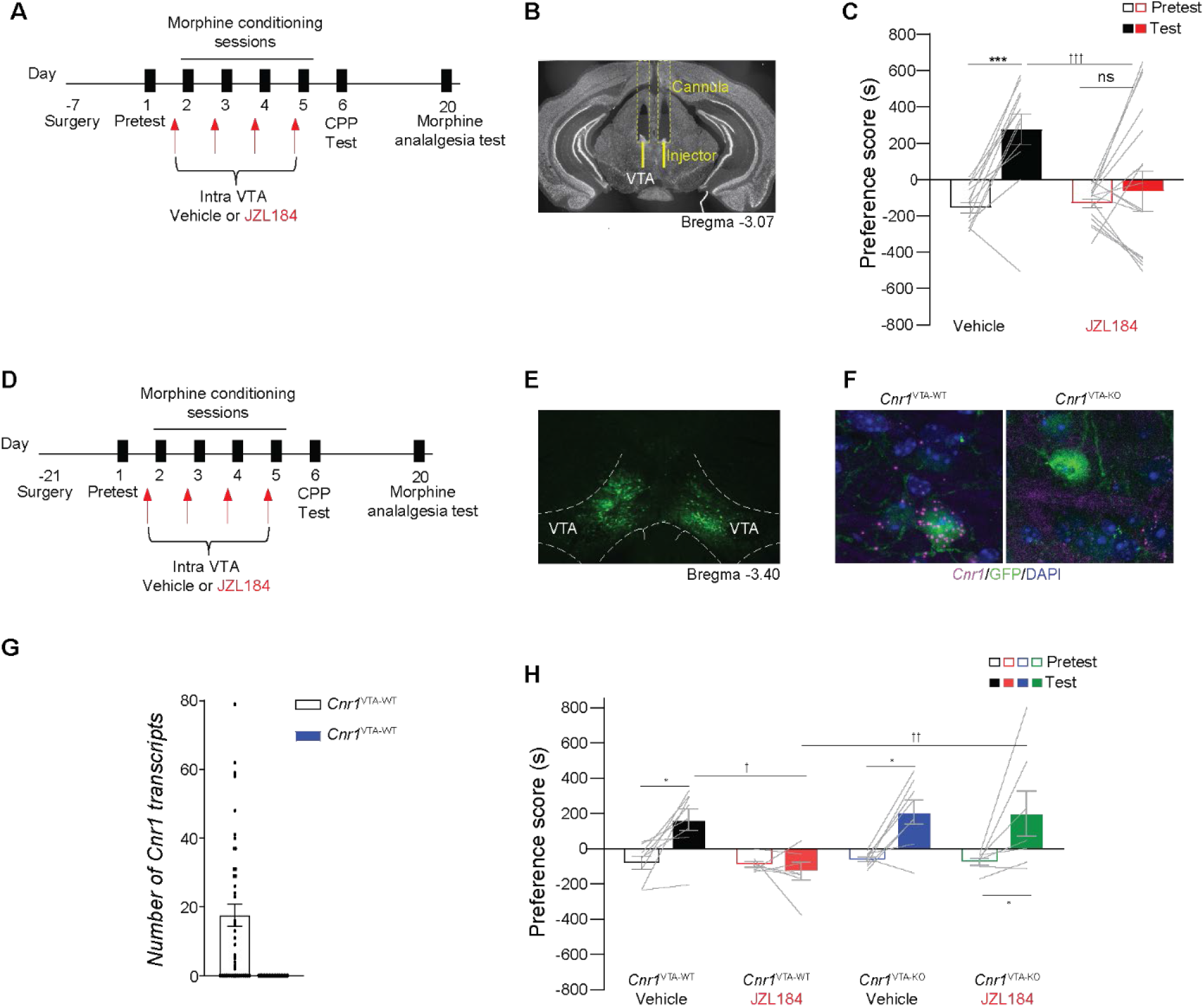
Intra-VTA JZL184 attenuates morphine preference but does not alter morphine analgesia. (**A**) Experimental timeline of surgery for guide cannula placement and intra-VTA infusion of JZL184 during morphine CPP, followed by tail-flick test to measure morphine analgesia. (**B**) Exemplar image of guide cannula placement in the VTA. (**C**) Intra-VTA infusion of JZL184 attenuates morphine preference (Two-way ANOVA, significant interaction (treatment x day), F_1,54_ =5.144, P=0.0274; Bonferroni post hoc test: Veh: Test vs Pretest, P=0.0051**, JZL184: Test vs Pretest no significance, P>0.999 Test: Veh vs JZL184 P=0.018*, Veh n=12, JZL184 n=17). (**D**) Experimental timeline for surgery, morphine CPP and morphine TFT. (**E**) Representative image of AAV2-Cre-GFP expression in the VTA (VTA=Ventral tegmental area). (**F**) Representative image of RNAscope in situ hybridization showing *Cnr1* mRNA expression (magenta) and GFP-tagged cells (green) in the VTA of *Cnr1* ^VTA-WT^ (left panel) and *Cnr1* ^VTA-KO^ (right panel) mice (DAPI= blue). (**G**) Quantification of *Cnr1* puncta in *Cnr1*^VTA-WT^ and *Cnr1*^VTA-KO^ mice (Ttest, t(_70_) = 4.508, P<0.001***, *Cnr1*^VTA-WT^ n=3 (42 cells in total), *Cnr1*^VTA-KO^ n=3 (25 cells in total). (**H**) Focal knockout of CB1Rs in the VTA counteract JZL184’s morphine blunted response (Three-way ANOVA, significant interaction (days x genotype), F_1,26_ = 4.530, P<0.043*; Bonferroni post hoc test: *Cnr1* ^VTA-WT^ Veh: Test vs Pretest, P=0.044*, *Cnr1* ^VTA-WT^ JZL184: Test vs Pretest no significance, P>0.999, *Cnr1* ^VTA-KO^ Veh: Test vs Pretest, P=0.020*, *Cnr1* ^VTA-KO^ JZL184: Test vs Pretest, P=0.031*, Test: *Cnr1* ^VTA-WT^ Veh vs *Cnr1* ^VTA-WT^ JZL184, P=0.013†, Test: *Cnr1* ^VTA-WT^ JZL184 vs *Cnr1* ^VTA-KO^ JZL184, P=0.005††, *Cnr1* ^VTA-WT^ Veh n=8, *Cnr1* ^VTA-WT^ JZL184 n=7, *Cnr1* ^VTA-KO^ Veh n=8, *Cnr1* ^VTA-KO^ JZL184 n=7).

### Knockout of VTA CB1R blunts JZL184’s effects on morphine CPP

Next, to test whether CB1R’s within the VTA are responsible for JZL184’s effects on morphine CPP, we generated focal knockout of CB1Rs in the VTA using CB1R gene, *Cnr1* floxed mice. The VTA of *Cnr1* wildtype (*Cnr1*^WT^) or *Cnr1* floxed mice (*Cnr1*^fl/fl^) were bilaterally microinjected with AAV2-Cre-GFP to generate *Cnr1*^VTA-WT^ and *Cnr1*^VTA-KO^ mice (**Figure 4D-F, Supplementary Figure 4F**). To validate the efficiency of *Cnr1* knockout in virus infected cells (**Figure 4E**) we performed dual GFP immunofluorescence and *Cnr1* RNAscope in situ hybridization (**Figure 4F**). Quantification of *Cnr1* puncta in *Cnr1*^VTA-WT^ and *Cnr1*^VTA-KO^ mice validated the viral strategy and confirmed the deletion of *Cnr1* mRNA in GFP-positive cells (**Figure 4G** RNAscope images; quantification). *Cnr1*^VTA-WT^ and *Cnr1*^VTA-KO^ mice were subjected to morphine CPP receiving systemic JZL184 pretreatment before each morphine conditioning session (**Figure 4D**). As expected, control *Cnr1*^VTA-WT^ animals with intact CB1Rs pretreated with vehicle acquired a preference for the morphine chamber, while JZL184 pretreated mice exhibited a blunted response (**Figure 4H**). The blunted response was reversed in *Cnr1*^VTA-KO^ mice treated with JZL184 with these animals acquiring a preference for the morphine-paired chamber, demonstrating that VTA CB1Rs are necessary for JZL184’s effect. Interestingly knockout of CB1Rs in the VTA did not affect morphine preference (*Cnr1*^VTA-KO^ treated with vehicle).

The analgesic response to morphine was not affected by the genotype (**Supplementary Figure 4G**). JZL184 pretreatment tends to decrease the ED_50_ values in both genotypes, when compared with their genotype homologous counterparts treated with vehicle, suggesting that deletion of VTA-CB1R’s within the VTA does not alter the morphine analgesic response.

### Pharmacologically inhibiting MAGL blunts nucleus accumbens (NAc) neural activity and DA neurotransmission

The VTA plays a critical role in opioid reward (32, 33) and reinforcement (34) primarily through activation of dopaminergic afferents projecting to the NAc that increase NAc activity (35) via DA neurotransmission. Given this, and our behavioral data indicating that enhancing 2-AG blunts opioid reward, we hypothesized that enhancing 2-AG may be preventing the increase in NAc activity and DA release associated with opioid reward. To measure neural activity, we used the genetically encoded Ca^2+^ indicator GCaMP6s alongside fiber photometry (FP) to record *in vivo* Ca^2+^signals as a proxy for neural activity (36–38). A virus for GCaMP6s (AAV1-Syn-GCaMP6s-WPRE.SV40) was injected unilaterally into the NAc and a fiber optic cannula was implanted above the injection site to deliver 470 nm excitation light and record changes in fluorescence (**Figure 5A-B, Supplementary Figure 5A**). Mice were tested in morphine CPP with vehicle or JZL184 pretreatment during morphine conditioning (as described for **Figure 1B**) and fiber photometry recordings of NAc activity were obtained during the CPP test session (Day 6; **Figure 5C**). Vehicle-treated mice showed an increase in NAc GCaMP6s signal as they approached the morphine-paired chamber, while conversely the signal decreased when approaching the saline-paired chamber (**Figure 5D, 5F**; **Supplementary Figure 5B, 5D**). This difference in signal persisted as mice entered the morphine chamber compared to the saline chamber (**Supplementary Figure 5C, 5D, 5F, 5H**). In contrast, no change in GCaMP6s signal was seen as mice pretreated with JZL184 approached (**Figure 5E, 5F)** or entered (**Supplementary Figure 5E, 5G, 5H**) either chamber. Thus, opioid place preference is associated with an elevation in NAc activity as mice approach and enter the opioid-associated chamber and this activity is fully abolished by JZL184 pretreatment during drug conditioning.

**Figure 5.**
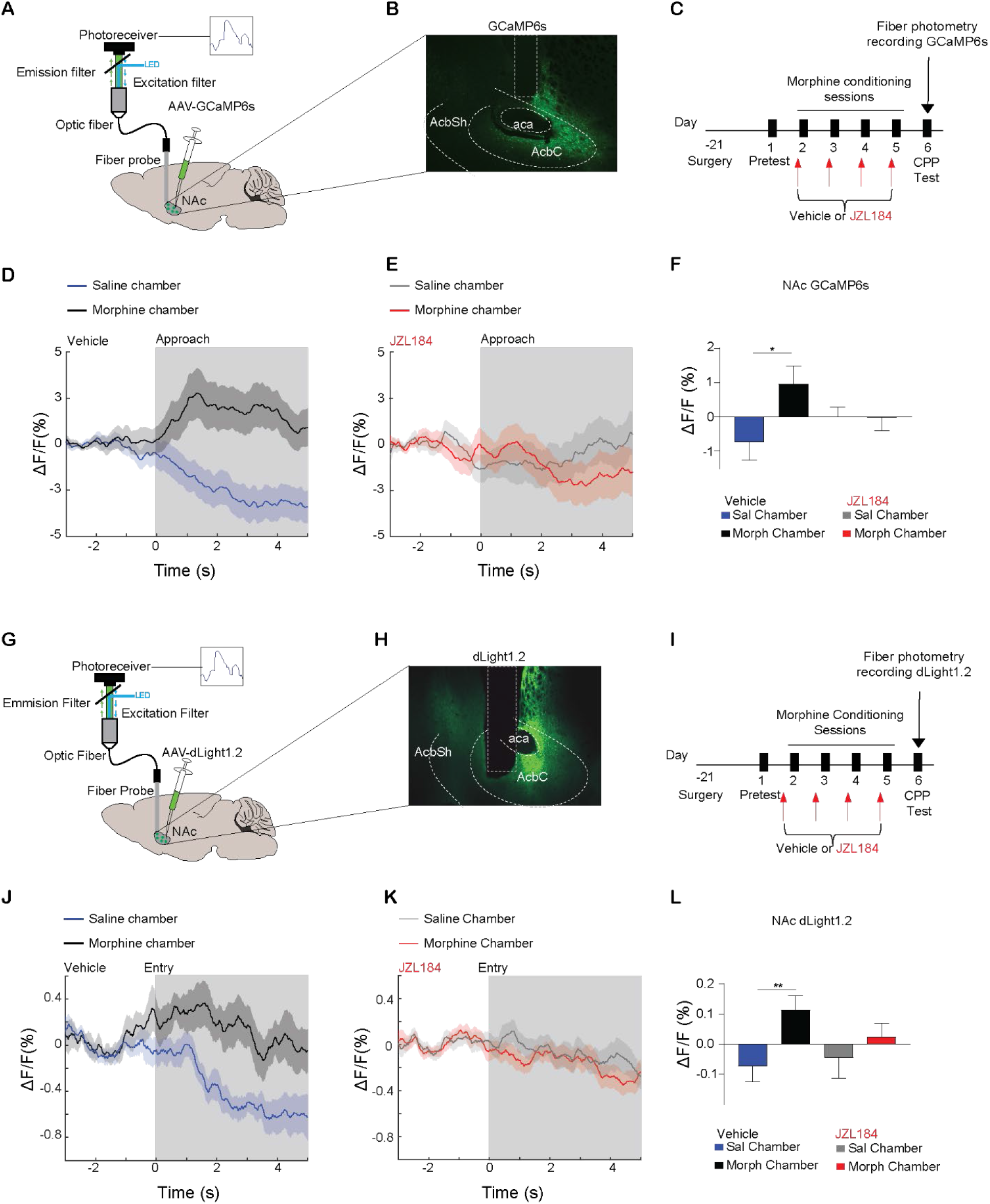
JZL184 pretreatment attenuates NAc neural activity and DA dynamics time-locked to the morphine-paired chamber. (**A**) Brain schematic with an optic fiber implanted into the NAc expressing GCaMP6s and recording apparatus. (**B**) Representative image of GCaMP6s expression in the NAc (aca=anterior commissure, AcbC=NAc core, AcbSh=NAc shell). (**C**) Experimental timeline for surgery, morphine CPP and fiber photometry recording of calcium (GCaMP6s) on CPP preference test session on Day 6. (**D**) A representative trace of the mean ΔF/F (+/– SEM) photometry trace of NAc GCaMP6s fluorescence in vehicle pretreated mice time-locked to the exit from saline-paired chamber approaching the morphine-paired chamber (black) or exit from morphine chamber approaching the saline chamber (blue). (**E)** A representative trace of the mean ΔF/F (+/– SEM) photometry trace of NAc GCaMP6s fluorescence in JZL184 pretreated mice time-locked to the exit from saline-paired chamber approaching the morphine-paired chamber (red) or exit from morphine chamber approaching the saline chamber (grey). (**F**) Vehicle but not JZL184 treated mice exhibit significantly higher ΔF/F when approaching the morphine-compared to the saline-paired chamber (linear mixed effects model, **p=0.0047, Vehicle, number of approaches = 191, n = 6; p=0.974 JZL184, number of approaches = 284, n = 8). (**G**) Brain schematic with an optic fiber implanted into the NAc expressing dlight1.2 and recording apparatus. (**H**) Representative image of dLight1.2 expression in the NAc (aca=anterior commissure, AcbC=NAc core, AcbSh=NAc shell). **(I**) Experimental timeline for surgery, morphine CPP and fiber photometry recording of DA (dligh1.2) on CPP preference test session on Day 6. (**J**) A representative trace of the mean ΔF/F (+/– SEM) photometry trace of NAc dlight1.2 fluorescence in vehicle pretreated mice time-locked to entry into the morphine-paired chamber (black) or saline-paired chamber (blue). (**K**) A representative trace of the mean ΔF/F (+/– SEM) photometry trace of NAc dlight1.2 fluorescence in JZL184 pretreated mice time-locked to the entry into the morphine-paired (red) or saline-paired chamber (grey). (**L**) Vehicle but not JZL184 pretreated mice exhibit significantly higher NAc ΔF/F when entering the morphine-paired compared to the saline-paired chamber (linear mixed effects model, **p=0.007, Vehicle, number of entries = 575, n = 16; p=0.369 JZL184, number of entries = 448, n = 12). Error bars ± SEM.

Next, we used FP to examine DA levels in the NAc using the genetically encoded fluorescent DA sensor, dlight1.2 (46). We hypothesized that JZL184 would decrease DA release in the NAc given that it abolished reward-related NAc activity (**Figure 5D-F**). Mice were injected unilaterally with a viral vector for dlight1.2 (AAV5-hSyn-dLight1.2) into the NAc and implanted with a fiber optic cannula above the injection site to deliver 470 nm excitation light and record changes in fluorescence (**Figure 5G, 5H, Supplementary Figure 6A**). Mice were tested in morphine CPP with vehicle or JZL184 pretreatment during conditioning (as described in Figure 1B) and FP recordings of DA activity were obtained during the CPP test session (Day 6; **Figure 5I**). Vehicle treated mice showed an increase in dlight1.2 signal when they entered the morphine-paired chamber while the signal decreased as they entered the saline-paired chamber (**Figure 5J, 5L, Supplementary Figure 6B).** This difference in dlight1.2 signal during entry was blunted in JZL184 pretreated mice (**Figure 5K, 5L, Supplementary Figure 6C**). There was no change in signal during approach behavior in vehicle or JZL184 pretreated mice (**Supplementary Figure 6D-F**). These results are consistent with our hypothesis that elevating 2-AG levels blunts opioid-induced dopaminergic transmission and neural activity within the NAc.

## Discussion

In this study we demonstrate that systemic or intra-VTA MAGL inhibition with JZL184 (that selectively elevates 2-AG; (39)) attenuates the rewarding effects of opioids through CB1Rs with no effect on analgesia. In contrast, enhancing AEA has no effect on opioid reward. Our *in vivo* calcium and DA detection studies indicate that enhancing 2-AG levels diminishes opioid reward-related NAc neural activity and DA transmission (**Figure 6**). These studies corroborate and expand upon previous pharmacological studies demonstrating a dampening effect of augmented 2-AG of opioid reward (15, 40).

**Figure 6.**
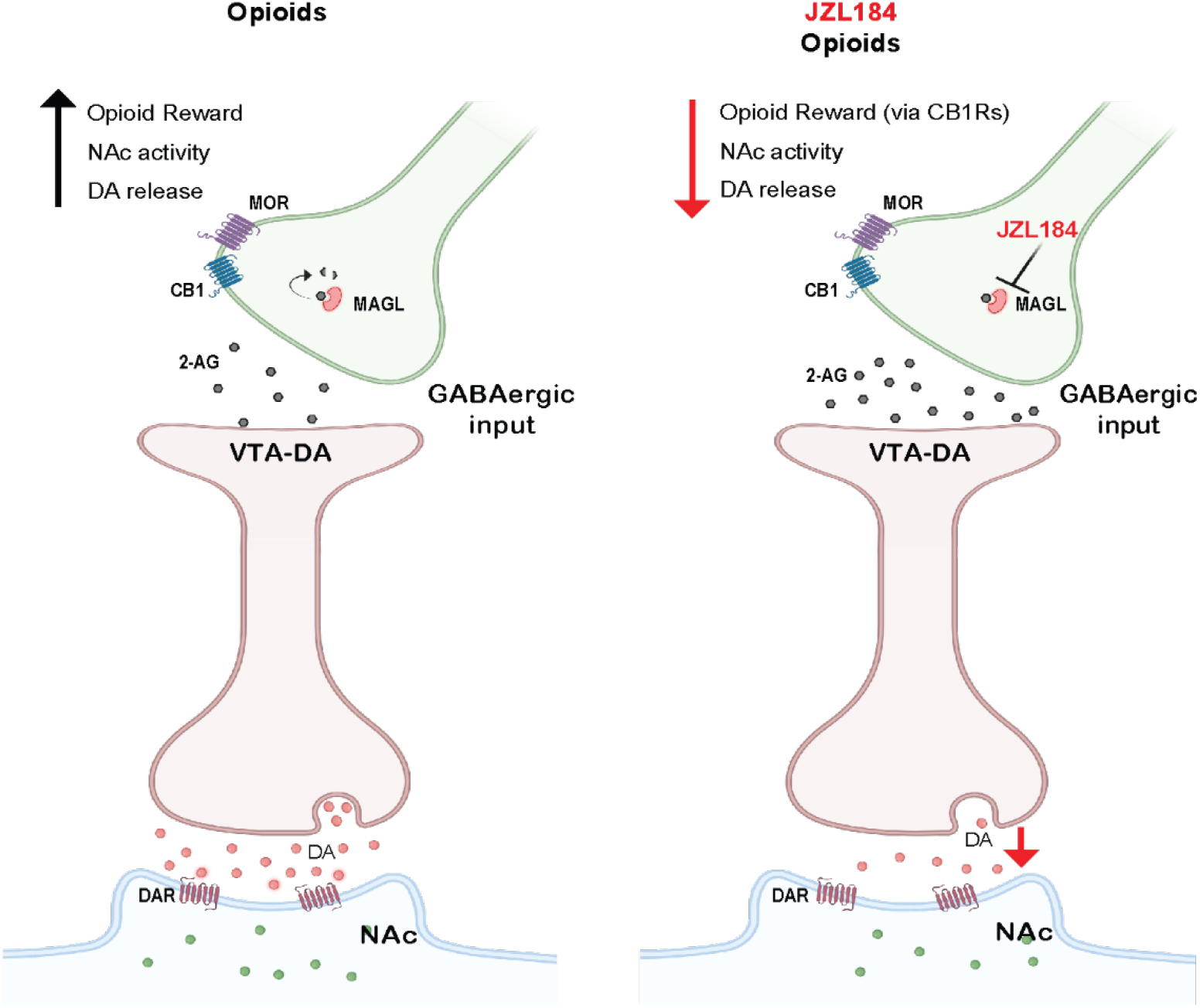
Morphine reward is associated with increase in NAc activity and dopamine neurotransmission that is blunted following 2-AG elevation in the VTA. Left, Opioid reward occurs via increase of NAc activity and DA levels. Right, Elevation of 2-AG in the VTA through inhibition of MAGL with JZL184 during opioid conditioning, blunts reward, NAc activity and DA levels. Ca^2+^ = calcium, DA=dopamine, DAR = dopamine receptor, CB1= cannabinoid receptor 1, MOR= µ-opioid receptor, 2-AG= 2-Arachidonoylglycerol.

The effect of enhancing 2-AG on attenuating opioid reward was unexpected for a few reasons. First, it has been established that exogenous cannabinoids such as THC, mediate reward and reinforcing behaviors through the disinhibition of VTA DA neurons, leading to enhanced DA transmission in the NAc (41, 42). In addition, THC has been shown to enhance the rewarding properties of opioids (43–45). Furthermore, antagonizing CB1R leads to decreased reward behaviors and DA release induced by cocaine, amphetamine, nicotine, and alcohol (46, 47), as well as impaired morphine analgesia (48). Finally, acutely, and repeatedly increasing 2-AG levels via MAGL inhibition has been shown to increase cue-evoked reward seeking behavior and enhanced DA levels in NAc (18, 49, 50). However, recently, a study finds that enhancing 2-AG levels can attenuate operant intracranial self-stimulation of the medial forebrain bundle, which is known to activate reward pathways (51).

In our study, JZL184 was administered at a relatively low dose (10 mg/kg) with daily administration over a 4-day period, likely leading to a protracted mild enhancement of 2-AG levels, as compared to the acute effects of high doses (18-40 mg/kg) of JZL184 previously reported for brain stimulation-based reward behavior (18, 49). In addition, this repeated low dosing schema has previously been shown not to affect CB1R expression or function with regards to antinociceptive properties of the MAGL inhibitor (52–54). Conversely, repeated high dosing of JZL184 (>16 mg/kg) led to functional downregulation of CB1R (54, 55). However, repetitive MAGL inhibition with low dose JZL184 (8 mg/kg, i.p.) was also recently shown to increase NAc DA levels and facilitate sucrose-based reward (50), highlighting the notion that opioid reward-related behavior may be uniquely regulated by the eCB system. This is further supported by our data of the ability of repeated JZL184 to abolish morphine CPP-associated NAc DA levels versus increase in dopamine associated with brain stimulation and sucrose reward (18, 50). While there is a notable gap in the literature on the impact of elevating the endogenous cannabinoids specifically on opioid reward-related behaviors, a previous study with a dual FAAH-MAGL inhibitor (SA-57) was shown to attenuate heroin self-administration (15), with no effect of a FAAH inhibitor (56), pointing to the contribution of MAGL inhibition on opioid seeking.

The selective attenuating effects of 2-AG on opioid reward, in our CPP and self-administration paradigms appear to be specific as our studies indicate that AEA does not alter opioid reward, as measured by CPP. Of note, of these CB1R agonists, 2-AG has the highest apparent affinity and efficacy for the cannabinoid receptor (57, 58). It will be interesting in the future to test whether the relative efficacy of a broader range of CB1R agonists on opioid reward behavior as well as NAc neural activity is dependent on distinct pharmacological properties.

Opioids are thought to mediate rewarding effects by acting on presynaptic MORs on GABAergic inputs onto VTA DA neurons (59, 60). In this model, presynaptic MOR activation disinhibits VTA DA neurons, enabling DA release and increased NAc activity. The rostromedial tegmental area (RMTg), also known as the tail of the VTA provides one of the major sources of morphine-sensitive inhibitory inputs to the VTA (61, 62), mainly to dopaminergic neurons (63), and optogenetic or chemogenetic behavioral studies have strongly implicated this projection in opioid reward (64–66). CB1Rs are also present on GABAergic inputs to the VTA, including the RMTg (67). We demonstrated that CB1-VTA-KO counteracted the effects of JZL184 on blunted morphine response. Therefore, CB1 within the RMTg might be the site of action of JZL184 on morphine reward, suggesting a possible endocannabinoid/opioid crosstalk. However, CB1Rs are also expressed among different neurons of the VTA (68, 69), including VTA Dopaminergic (68), glutamatergic (69) and GABAergic neurons (69, 70). Currently, it remains unclear if the effects of JZL184 are dependent on CB1Rs within one specific population of neurons in the VTA. Further studies are required to determine and validate the assumption that the effects of JZL184 depend on RMTg input to the VTA. Notably, endocannabinoid activation specific within opioid-sensitive inputs to the VTA may provide an explanation for the specific effects of JZL184 on opioid reward but not on opioid analgesia, which is likely mediated further downstream at dopaminergic synapses in the NAc and other pain-sensitive brain regions, respectively.

In summary, we have identified a new functional interaction between the opioid and eCB systems, by which protracted elevation of the eCB 2-AG leads to attenuated opioid-mediated reward. These findings highlight an unexpected role of 2-AG in regulating reward processes that is distinct from other cannabinoid ligands. In addition, the present findings provide a compelling rationale for developing a new class of adjunctive endocannabinoid-based treatments that could dissociate the rewarding and analgesic properties of opioids.

## Material and Methods

### Animals

C57BL/6 male mice (Charles River Laboratories, Wilmington, MA) were 8-10 weeks of age at the start of the experiments. Mice were provided with food and water ad libitum. Animals were kept on a 12-hr light/dark cycle (from 7 A.M. to 7 P.M.).

### Drugs and inhibitors

Morphine was obtained through the National Institute on Drug Abuse following the Ordering Guidelines for Research Chemicals and Controlled Substances. Cocaine HCl and Oxycodone HCl were purchased from Sigma (St. Louis, MO). The FAAH inhibitor PF-3485, and the MAGL inhibitor JZL184 were obtained from Cayman Chemical. PF-3845 and JZL184 were dissolved in DMSO and diluted to their respective final concentrations in a solution of DMSO, Tween 80 and Saline at a ratio of 10:10:80.

### Conditioned place preference (CPP)

As previously published opioid (31) conditioned place preference was performed using a three-chamber (black, white and a center gray chamber) apparatus (Med Associates, Inc). We use a 6-day protocol with four days of conditioning. Briefly, on day 1, mice were placed in the central gray chamber for one minute of habituation followed by free exploration in the three chambers (pre-test). Times spent in the black and white chamber were recorded. During conditioning days, in the morning mice were injected with saline (i.p.; 0.01 ml/g body weight) and confined to the chamber they preferred during pretest (20 minutes for morphine, and 30 mins for oxycodone). In the afternoon mice were injected with morphine (10 mg/kg, i.p.), or oxycodone (1 mg/kg, i.p.) and confined for 20 mins (morphine) or 30 mins (oxycodone). For pharmacological experiments mice were pretreated with JZL184 (10 mg/kg) and PF-3845 (10 mg/kg) or vehicle (10% DMSO, 10% Tween 80 in saline) were administered two hours prior to morphine treatment (25, 26). Preference score was calculated as time spent in the cocaine-paired chamber minus time spent in the saline-paired chamber. Mice are defined as having acquired morphine conditioning place preference when the preference score during the test session (Day 6) is significantly higher than the preference score on the preconditioning test (Day 1). Locomotor activity (based on total beam breaks) was obtained while mice were in the CPP chambers.

### Jugular Catheterization Surgery

Prior to surgery, mice were anesthetized with 80 mg/kg ketamine and 12 mg/kg xylazine. An indwelling silastic catheter was placed into the right jugular vein and sutured in place. The catheter was then threaded subcutaneously over the shoulder blade and was routed to a mesh back mount platform (Instech Laboratories, Inc.) that secured the placement. Catheters were flushed daily with 0.1 ml of an antibiotic (Timentin, 0.93 mg/ml) dissolved in heparinized saline.

### Operant food training

Three days following catheterization, mice were trained to perform an operant response for sucrose pellets. The mice were placed in operant chambers (Med Associates) and trained to press a lever to receive a sucrose pellet. Mice performed two consecutive days of FR1 responding. A compound cue stimulus consisting of a cue light above the active lever, a 2900-Hz tone, and house light off was concurrent with each pellet administration, followed by an additional 8 second time-out when responding had no programmed consequences and the house light remained off. Mice were allowed to self-administer a maximum of 50 pellets per self-administration session. During the food training phase, mice were food restricted to >90% of their free-feeding weight. Mice returned to ad libitum food access 5 days following the start of oxycodone self-administration.

### Oxycodone-Self Administration

Mice were tested for oxycodone self-administration behavior in 2-hour sessions in the same chamber used for sucrose pellet self-administration. Animals were randomly assigned to receive either JZL184 or vehicle (10% DMSO, 10% Tween 80 in saline or saline alone; there was no significant difference in behavior between these two controls groups and thus they were combined) i.p. injections daily, 2-hours prior to the start of each of the oxycodone self-administration sessions (wheels and levers). Mice were first tested on three consecutive days of oxycodone self-administration using a wheel. Mice were trained to spin a wheel (Med Associates) quarter of a turn that delivered an intravenous oxycodone injection (0.25mg/kg/infusion). During testing, responding on the wheel delivered an intravenous oxycodone infusion (0.25mg/kg/infusion), paired with the same compound cue under the same schedule as the food training, followed by an 8 s time-out when responding had no programmed consequences. Following three days of wheels, mice were transitioned from a wheel back to a lever, which they had previously learned to press during the operant food training phase. Press of an active lever delivered an intravenous oxycodone injection (0.25mg/kg/infusion) whereas press of an inactive did not have any consequences. Mice were evaluated for oxycodone self-administration on an FR1 schedule for an additional seven days. The percent of correct choices during the lever press sessions was calculated using the formula (number of active lever press)/(number of active lever press **+** number of inactive lever press)**\***100.

### Hot Plate Assay

Mice were tested for analgesia using the Hot/Cold Plate apparatus (UGO Basile, S.R.L.). First, mice were tested for their baseline response, animals were collocated on the platform of the hot plate set at 55 °C. The time(s) elapsing to the first pain response (licking or jumping) was scored. Then, mice were injected with JZL184 (10 mg/kg, i.p.) or vehicle, and two hour later mice were tested for morphine analgesia with a cumulated dose-response curve (0.3 mg/kg – 10 mg/kg), using the hot plate. Mice were tested 30 min after each dose was administered. A maximal latency of 30 sec was used to minimize any tissue damage. Results were determined as %MPE ([latency after drug − baseline latency]/[30 − baseline latency]*100).

### Tail Flick Test

Mice were tested for analgesia using the Ugo Basile 37360 tail flick device. For the baseline test, mice were wrapped and the lower 1/3 of the tail was placed over the sensor of the device which emitted radiant heat. The animals’ response to the heat is a flick of the tail. The latency of the tail flick was obtained as a measure of the baseline response with a maximal latency of 10 seconds to minimize tissue damage. After recording the value, mice were injected with JZL184 or PF-3845, followed by the pretreatment waiting time of two hours. Then the mice received the lower dose of morphine or oxycodone injection followed by a 30-minute waiting time. Mice were tested for analgesia (the same way the baseline was performed in terms of tail flick), and the maximum time the tail is exposed to the tail flick device is 10 seconds to avoid severe burns. Immediately after the first dose test, mice were injected with the second dose and 30 min later the tail flick test was repeated. This was performed with 0.1, 0.3, 1.0, 3.0 and 10 mg/kg of morphine or 0.1, 0.3, 1.0, 3.0 and 5 mg/kg of oxycodone. Results were calculated as the percentage of maximum possible effect (% MPE) as follows: [(latency after drug – baseline latency)/(10 – baseline latency) × 100].

### Elevated Plus Maze (EPM)

EPM testing was performed as previously reported (71). Briefly mice were placed in the center of an elevated cross-shaped maze. Behavior was video recorded for 5 mins and analyzed using the any-maze software. Data is reported as the percent of time spent in the open arms which was calculated as: (time in open arms (s)/ total time (s)) *100.

### Tail Suspension Test (TST)

TST was performed as previously published (72). Briefly, TST was performed by suspending a mouse 30 cm from the floor using a 17 cm-long adhesive tape which was secured to the tail 2 cm from the tip. Each mouse was video recorded for 6 mins, and time spent immobile was hand-scored by an experimenter blinded to the treatments using the computer-assisted software CowLog.

### Intracranial Surgery

For the delivery of JZL184 into the VTA, guide cannulas were implanted bilaterally in adult mice as previously published (73). The coordinates used to inject JZL184 or vehicle, were AP= 0.4 to 0.7mm, ML= -3.1 to -3.3mm, and DV= 4.2 to 4.4mm. A total of 5ug/ul, 200nl/side/min of JZL184 was bilaterally microinjected 30 min before each morphine conditioning sessions (74).

For the in vivo fiber photometry experiments surgical procedures were conducted as previously published (75). A viral vector expressing GCaMP6s (AAV1-Syn-GCaMP6s-WPRE.SV40, from Addgene), or dLight1.2 (AAV5-hSyn-dLight1.2, from Adgene), was injected unilaterally into the NAc. The coordinates used to inject the virus were 1.0-1.2 mm AP relative to bregma, 0.75-0.85 mm ML and -4.5 to -4.75 DV (using Paxinos and Franklin’s the Mouse Brain in Stereotaxic Coordinates atlas as a reference). A total of 250 nl of GCaMP6s, or 600-800nl of dLight1.2 at a rate of 50 nl/min was injected unilaterally. Additionally, during the same surgery a 400 µm diameter optical fiber (Doric) was implanted approximately 0.2 mm dorsal of the virus injection site and was secured with Metabond. Following surgery, mice were housed with ad libitum access to food and water and were allowed a minimum of 21 days for viral expression before all experiments. After the completion of all the behavioral experiments, accurate virus injection was confirmed by immunohistochemistry; mice with improper injection or fiber placement were excluded from fiber photometry analyses.

For conditional knockout of CB1, AAV2-GFP-Cre (Vector Biolabs) were bilaterally injected into the VTA of *cnr1* wildtype *(Cnr1*^WT^) or *cnr1* floxed mice *(Cnr1*^fl/fl^) (*76*). The coordinates used to inject the virus were -3.1 mm AP relative to bregma, 0.4 mm ML and -4.4 DV (using Paxinos and Franklin’s the Mouse Brain in Stereotaxic Coordinates atlas as a reference). A total of 300 nl of AAV2-GFP-Cre, at a rate of 100 nl/min was injected bilaterally. Following surgery, mice were housed with ad libitum access to food and water and were allowed a minimum of 21 days for viral expression before all experiments. After the completion of all the behavioral experiments, accurate virus injection was confirmed by immunohistochemistry and focal knockout of CB1Rs within the VTA was confirmed with RNAscope. Mice with improper injection were excluded from statistical analyses.

### In vivo fiber photometry recording

*In vivo* fiber photometry was performed to measure calcium-dependent activity and DA levels during CPP behavior as previously described (36). Briefly, two to three days prior to behavioral testing, mice were habituated to a patch cord attached to the implanted optical fiber for at least 1 minute in the animal’s home cage. During morphine CPP test (Day 6), fiber photometry recordings were taken during the 20-minute CPP test. Additionally, the behavioral test was video recorded by a camera above the CPP apparatus. The fluorescent signal was normalized and transformed to ΔF/F by taking the median value of the overall trace, subtracting this median value from all data points and then dividing by the median. The NAc GCaMP6s and dlight1.2 fluorescent signals were time-locked to hand-scored morphine CPP behavior. GCaMP6s fluorescence was analyzed from 0 to 5 seconds from time of approach (i.e. timepoint when mouse leaves the opposite chamber) or entry (i.e. timepoint when mouse first enters the chamber) to the saline- or morphine-paired chamber.

### Histology

After completion of behavioral tests, animals were transcardially perfused with 0.1M phosphate buffer followed by 4% PFA, pH 7.4 and processed described previously (75). Nissl staining for guide cannula placements or GFP immunohistochemistry was performed for GCaMP6s, dlight1.2 viral placement.

### RNAscope

For the RNAscope, fixed brains were sectioned at 30 μm, and VTA sections were mounted on charged slides using 1x PBS, then allowed to air dry overnight. The next day, the mounted tissue was baked using the HybEZ oven (ACDbio) at 60°C for one hour. The sections were then acclimated for 10 seconds in near-boiling deionized water (dH2O) and incubated in 1X target retrieval buffer (at a temperature of 98-102 °C) for 5 minutes, followed by three washes using room temperature dH2O. The sections were then submerged in 100% ethanol for 3 minutes and allowed to air dry, followed by two washes with dH2O. All subsequent incubation periods were performed at 40°C and washes for 2 min each. The sections were then incubated in Protease III for 30 minutes, washed four times with dH2O, and then incubated with either a positive probe, negative probe, or *cnr1* probe for two hours. Slides were then washed in 1x wash buffer. Sections were incubated for 30 minutes with AMP1, AMP2 for 30 minutes, and AMP3 for 15 minutes, with one wash with 1x wash buffer performed in-between the previous incubation steps. Next, sections were incubated in HRP C1 for 15 minutes, followed by two washes with 1x wash buffer. Then sections were incubated with CY5 in TSA buffer for 30 minutes, followed by two washes and incubation with HRP-blocker for 15 minutes, with two subsequent washes. In order to investigate the co-localization of GFP-tagged virus with cacan1c mRNA, immunohistochemistry for GFP was conducted.

For imaging and quantification of RNAscope, images were taken as a z-Stack, with bidirectional X and 4 frame averaging. Filter used was the MBS 488/561/633. After acquisition at 40x and 20x magnification, images were processed for quantification using Fiji/ImageJ. One scan from the *Cnr1*^VTA-WT^ or *Cnr1*^VTA-KO^ was chosen and placed through automatic thresholding of the GFP channel. Using Analyze Particles, automatic region of interest (ROIs) was made around the soma of each GFP tagged neuron and added to the ROI manager. Using Find Maxima in ImageJ, a total number of puncta for the *Cnr1* probe were gathered within each ROI. Comparisons were made between control and experimental groups for *Cnr1* (CB1) mRNA transcripts in AAV2-CRE-GFP infected neurons.

### Statistical Analysis

Graph Pad Prism 8.0 was used for the analyses of the behavioral tests while MATLAB with custom scripts was used for the analyses of the in vivo fiber photometry recordings. For all the behavioral data a Two-way ANOVA statistical analysis was performed followed by a Bonferroni post-hoc test. For analysis of the fiber photometry recordings, MATLAB was used to perform a mixed linear effects model to compare average signal amplitude (ΔF/F) 0-5 seconds while approaching the chamber (GCaMP6s) or upon entry (dlight1.2) to the respective chambers, with the animal modeled as a random effect and CPP chamber (drug vs. saline) modeled as a fixed effect. Exact sample sizes are indicated in figure legends for each experiment. Statistical significance was defined as p value < 0.05.

### Study Approval

All procedures were conducted in accordance with the Weill Cornell Medicine Institutional Animal Care and Use Committee rules.

## Author Contributions

Experiments were conceptualized by AM-R, FSL, AMR, LAB and Y-XP. Methodology was designed by AM-R, FSL, AMR, JL, Y-XP, and CL. Reagents were obtained from VMP, KP. The investigation was performed by AM-R, RNF, CF, RY, JX and NM. Visualization of the project was by AM-R, FSL, AMR, LAB, Y-XP and JL. The main funding acquisition was obtained by FSL, AMR, JL, LAB, and Y-XP. The project administration and supervision was in charge of AM-R, FSL, and AMR. The writing of the original draft was performed by AM-R, FSL, AMR, JL and the writing-review & editing by AM-R, FSL, AMR, JL, Y-XP, and CL.

## Acknowledgements

We would like to thank Huda Akil, Alan Schatzberg, Jack Barchas, as well as other investigators in the Pritzker Neuropsychiatric Disorders Research Consortium for helpful discussions. We also extend our gratitude Dr. Joel Elmquist and Dr. Teppei Fujikawa from University of Texas Southwestern Medical Center for the generation of *Cnr1* floxed mouse. Additionally, we appreciate the contribution of Mathew Bredder, James Ryan and Viraj Nadkarni for help with fiber photometry analysis and Alexander Walsh for assistance with immunohistochemistry.

## Data and materials availability

All data are available in the main text or the supplementary materials. MATLAB code for fiber photometry data analysis is available upon request.

## Supplementary Materials

**Fig. S1.**
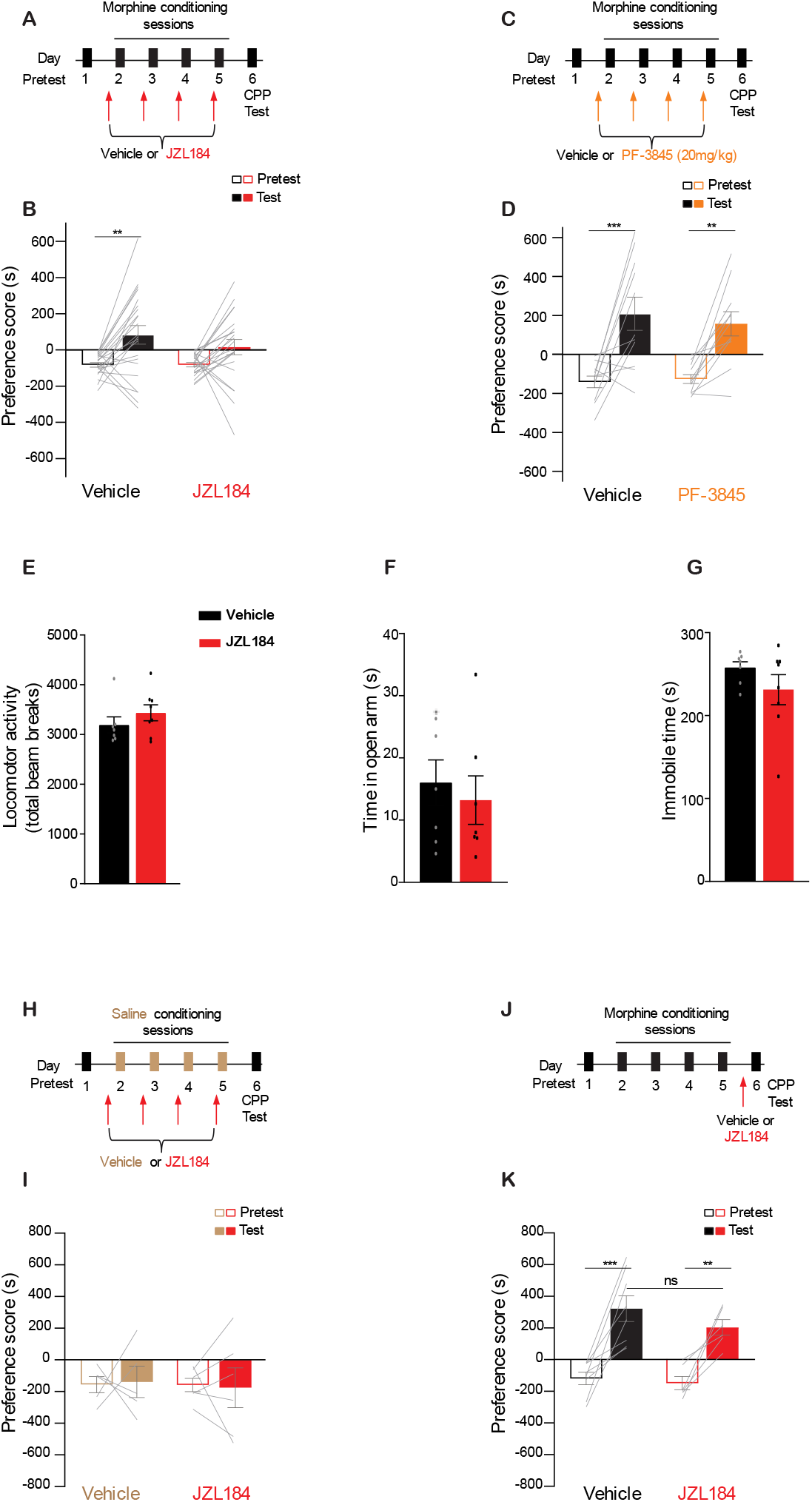
JZL184 attenuates morphine CPP in female mice, PF-3845 has no effect on morphine CPP in male mice, and JZL184 has no effect on the expression of morphine CPP, locomotion, anxiety-like behavior, and depressive-like behavior. **(A)** Timeline of behavioral protocol for morphine CPP and JZL184 (10mg/kg) systemic injections pretreatment in female mice. (**B**) Systemic injection of JZL184 (Two-way ANOVA, main effect of day, F_1,84_=0.95, P<0.0001; Bonferroni post hoc test: Veh: Test vs Pretest P=0.002**, JZL184: Test vs Pretest P=0.095, Veh n=22, JZL184 n=22) had no effect on morphine CPP. (**C**) Timeline of behavioral protocol for morphine CPP and PF-3845 (20mg/kg) systemic injections pretreatment. (**D**) Systemic injection of PF-3845 (20mg/kg) had no effect on morphine CPP (Two-way ANOVA, main effect of day, F_1,40_=31.79, P<0.0001; Bonferroni post hoc test: Veh: Test vs Pretest P=0.0002***, PF-3845: Test vs Pretest P=0.0019**, Veh n=11, PF-3845 n=11). (**E**) Locomotor activity on morphine CPP test day is not significantly different between vehicle and JZL184 pretreated mice. (**F-G**) Time in the open arm of the elevated plus maze (**F**) and immobile time in the tail suspension test (**G**) is not significantly different between vehicle and JZL184 pretreated mice. (**H**) Timeline of behavioral protocol for JZL184 on its own during CPP. (**I**) Systemic JZL184 on its own has no effect on conditioning place preference. (Veh n=5, JZL184 n=6). (**J**) Timeline of behavioral protocol for morphine CPP and systemic injection of JZL184 before the conditioning test. (**K**) Mice pretreated with systemic injection of JZL184 prior to expression test exhibited significantly higher morphine CPP similar to vehicle pretreated mice (Two-way ANOVA, main effect of day, F_1,24_=44.82, P=0.0001; Bonferroni post hoc test: Veh: Test vs Pretest P<0.001***, JZL184: Test vs Pretest P<0.0038**, Veh n=8, JZL184 n= 6). Error bars ± SEM.

**Fig. S2.**
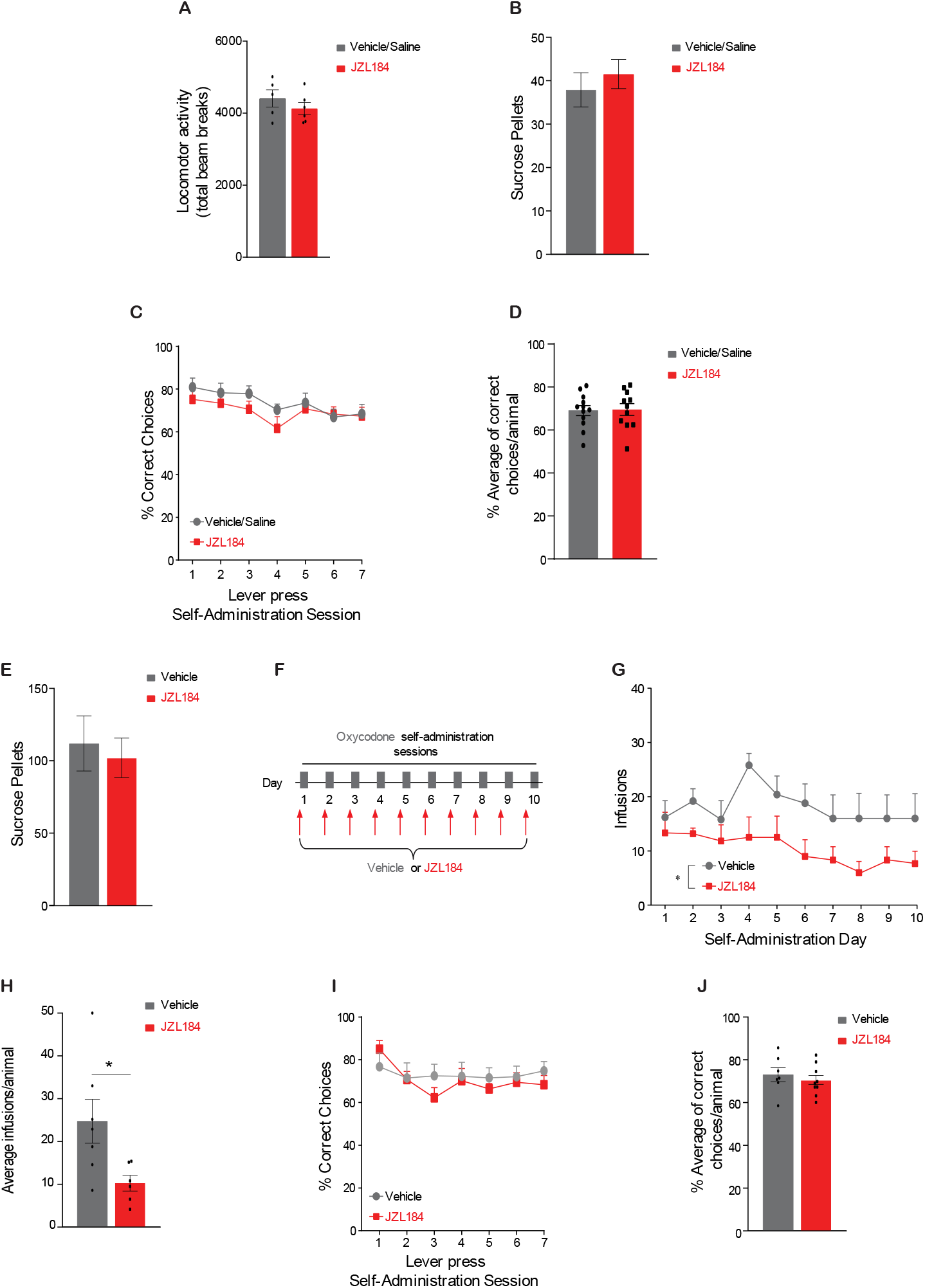
JZL184 pretreatment does not alter locomotor activity or the percentage of correct choices during oxycodone self-administration. (**A**) Locomotor activity on oxycodone CPP test day is not significantly different between vehicle and JZL184 pretreated mice. (**B**) Mice were trained in the operant self-administration apparatus with sucrose pellets, then they were assigned to receive JZL184 or Vehicle/Saline, no differences between the groups were detected during the self-administration of sucrose pellets. (**C**-**D**) Percentage of correct choices during oxycodone self-administration does not differ between JZL184 or Vehicle/Saline across days (**C**) or across animals (**D**). (**E**) Female mice were trained in the operant self-administration apparatus with sucrose pellets, then they were assigned to receive JZL184 or Vehicle/Saline, no differences between the groups were detected during the self-administration of sucrose pellets. (**F**) Timeline of behavioral protocol for oxycodone self-administration and JZL184 systemic injections pretreatment. (**G**) Systemic JZL184 exposure prior to oxycodone self-administration sessions, attenuated the intake of oxycodone (**G;** Two Way-RM ANOVA, main effect of JZL184 treatment, F_1,14_ =5.960, P=0.029*, main effect of days, F_2.482,34.75_ =11.01, P<0.0270*, **H;** Average of total infusions/animal (Ttest, t(_11_) = 2.477, P=0.031*, Veh n=7, JZL184 n=6). (**I**-**J**) Percentage of correct choices during oxycodone self-administration in females does not differ between JZL184 or Vehicle/Saline across days (**I**) or across animals (**J**). Error bars ± SEM

**Fig. S3.**
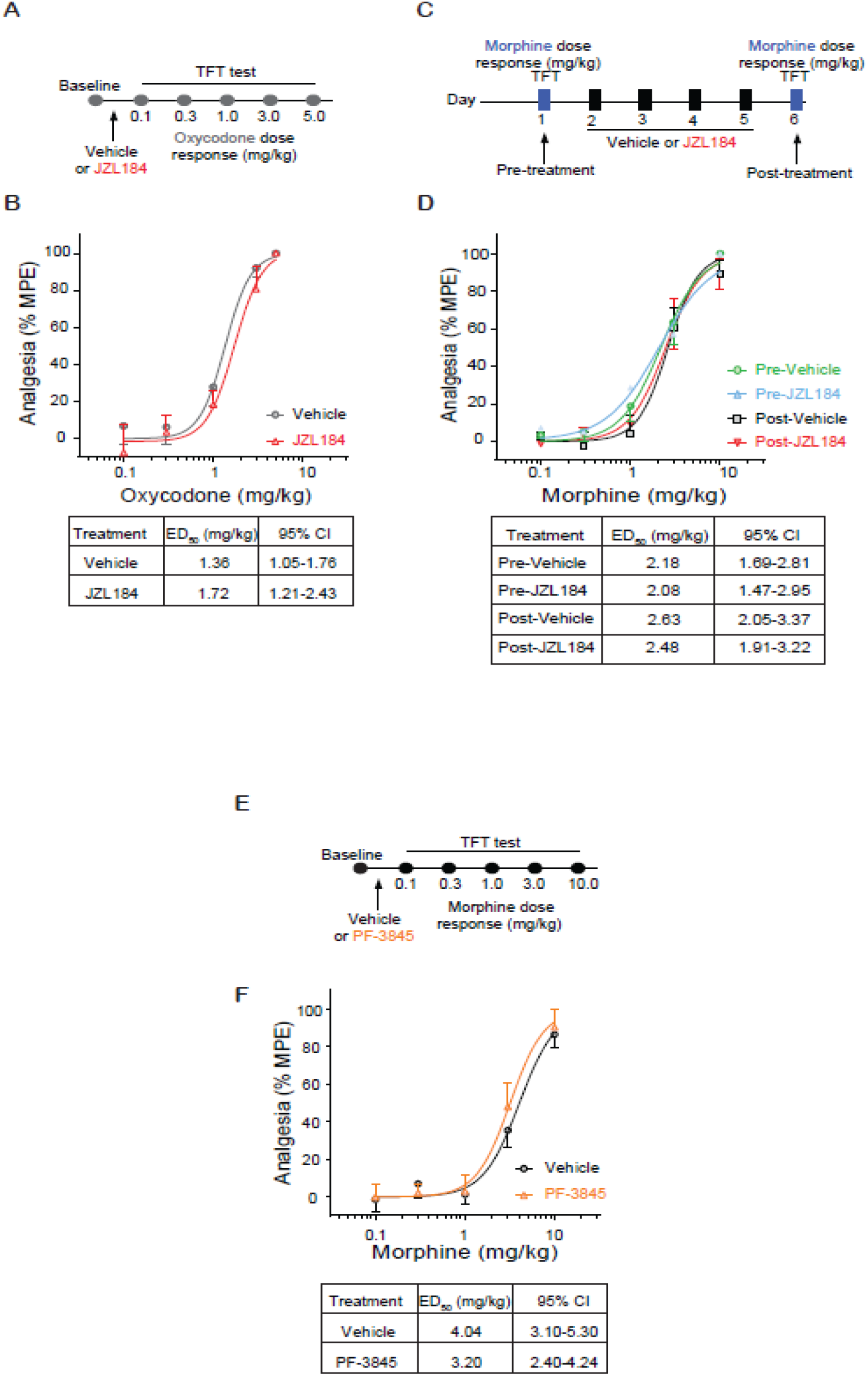
JZL184 and PF-3845 do not alter opioid analgesia. (**A**) Experimental timeline of acute systemic vehicle or JZL184 pretreatment prior to the oxycodone dose response in the tail-flick test. JZL184 pretreatment has no effect on oxycodone-induced analgesia (Two-way ANOVA, main effect of dose, F_4,55_= 53.44, P<0.001*** Veh n=7, JZL184 n=6). (**C**) Experimental timeline of tail-flick test to measure morphine analgesia pre and post repeated vehicle or JZL184 treatment. (**D**) Repeated JZL184 does not alter morphine analgesia (Two-way ANOVA, main effect of dose, F_4,130_ =112.6, P<0.001***, Veh n=7, JZL184 n=8). (**E**) Experimental timeline of acute systemic vehicle or PF-3845 pretreatment prior to the morphine dose response in the tail-flick test (**F**) The ED_50_ value of systemic PF-3845 pretreated animals were comparable to that in animals that received vehicle. (Two-way ANOVA, main effect of dose, F_4,80_ = 44.34, ***P<0.001; Veh n=11, PF-3845 n=7). Error bars ± SEM.

**Fig. S4.**
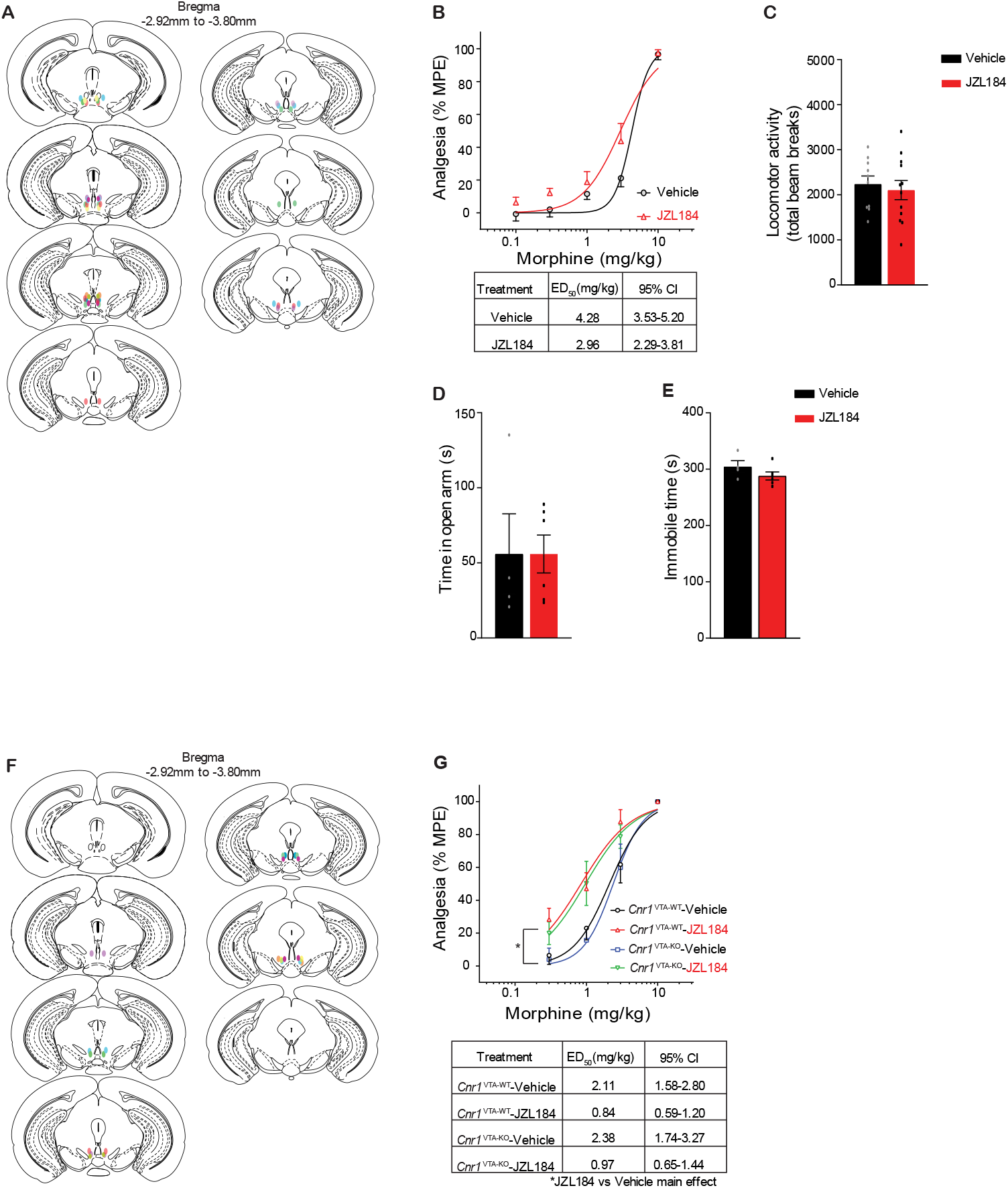
Brain placements and behavioral data for intra-VTA JZL or VTA-CB1 deletion during morphine CPP. (**A**) Coronal sections of JZL184 guide cannula placement in the VTA. Each animal received bilateral implantation. (**B**) Intra-VTA JZL184 does not alter morphine analgesia (Two-way ANOVA, main effect of dose, F_4,105_ = 107.7, P<0.001***, Veh n=11, JZL184 n=12). (B) Locomotor activity on morphine CPP test day is not significantly different between intra-VTA vehicle and JZL184 pretreated mice. (**D, E**) Time in the open arm of the elevated plus maze (**D**) and immobile time in the tail suspension test (**E**) is not significantly different between intra-VTA vehicle and JZL184 pretreated mice. (**F**) Coronal sections of AAV2-GFP-Cre expression in the VTA of Cnr1 floxed mice. (**G**) Focal knockout of CB1Rs in the VTA does not alter morphine analgesia effects (Three-way ANOVA, no genotype main effect or interaction, main effect of treatment JZL184 vs Veh F_1,60_ = 10.73, P<0.0019*, and significant interaction of Treatment (JZL184 or Veh) x dose, F_3,180_ = 3.601, P<0.015*; (*Cnr1* ^VTA-WT^ Veh n=9, *Cnr1* ^VTA-WT^ JZL184 n=9, *Cnr1* ^VTA-KO^ Veh n=7, *Cnr1* ^VTA-KO^ JZL184 n=11). Error bars ± SEM.

**Fig. S5.**
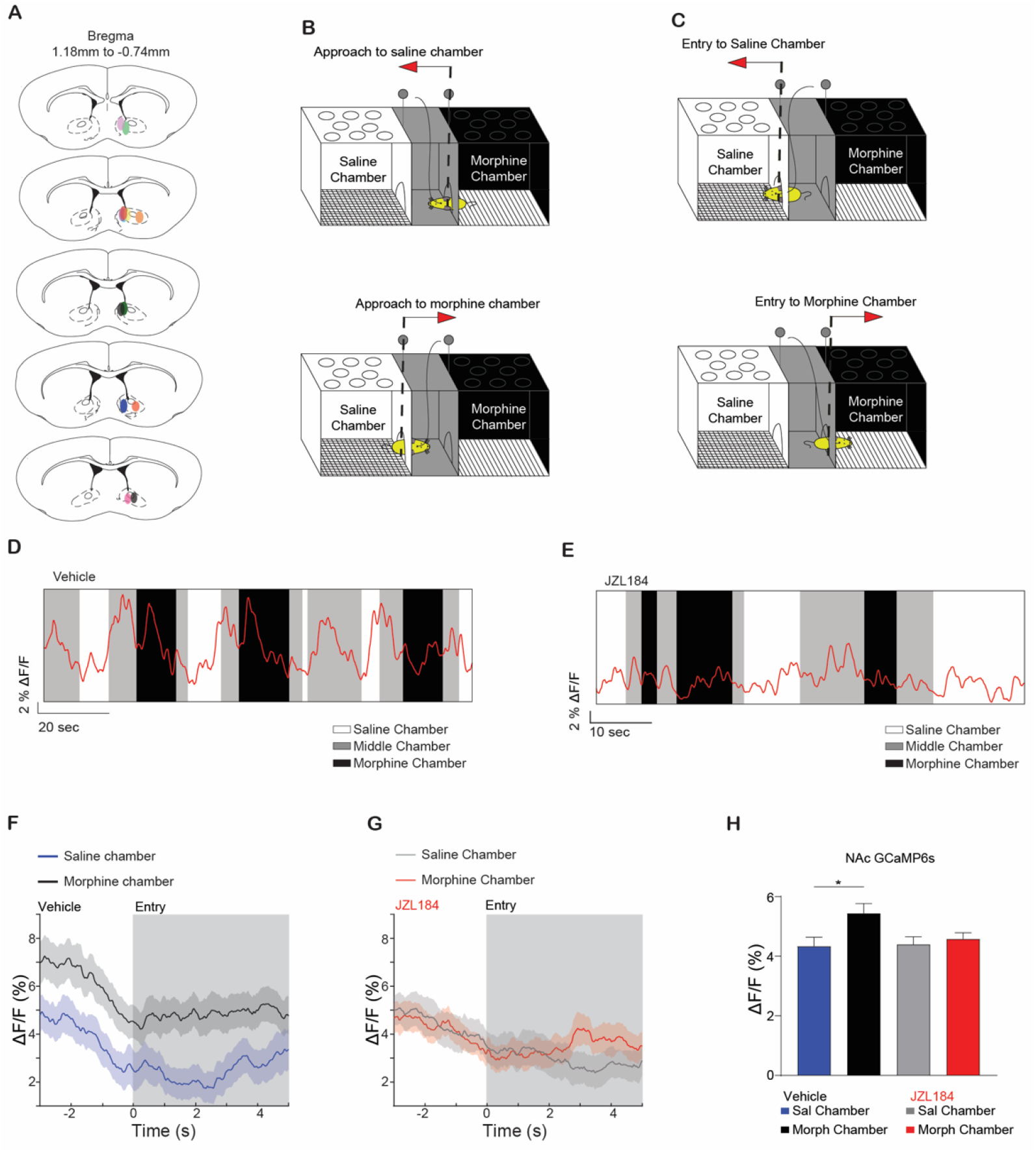
Brain placements and behavioral NAc GCaMP fiber photometry recordings. (**A**) Coronal sections of the fiber implant and GCaMP6s viral expression in the NAc. (**B**) Schematic of the CPP box depicting a mouse approaching the saline chamber (Top) or the morphine chamber (bottom). Dashed line represents the position in the CPP box where the fiber photometry GCaMP6s or dlight1.2 signal was time locked to the approach behavior. (**C**) Schematic of the CPP box depicting a mouse entering the saline chamber (Top) or the morphine chamber (bottom). Dashed line represents the entry point in the saline or morphine chamber of the CPP box to which the fiber photometry GCaMP6s or dlight1.2 signal was time locked. **(D-E)** Representative GCaMP6s traces collected during the CPP preference test day 6 in vehicle (**D**) or JZL184 (**E**) pretreated animals. (**F**) A representative trace of the mean ΔF/F (+/– SEM) NAc GCaMP6s fluorescence signal in vehicle pretreated mice time-locked to entry into the morphine-paired (black) or the saline-paired chamber (blue). (**G)** A representative trace of the mean ΔF/F (+/– SEM) NAc GCaMP6s fluorescence signal in JZL184 pretreated mice time-locked to entry into the morphine-paired (red) or saline-paired chamber (gray). (**H**) Vehicle but not JZL184 treated mice exhibit significantly higher ΔF/F when entering the morphine-compared to the saline-paired chamber (linear mixed effects model, Vehicle: *p=0.0160, number of entries = 248, n = 6; JZL184: p=0.5412, number of approaches = 437, n = 8).

**Fig. S6.**
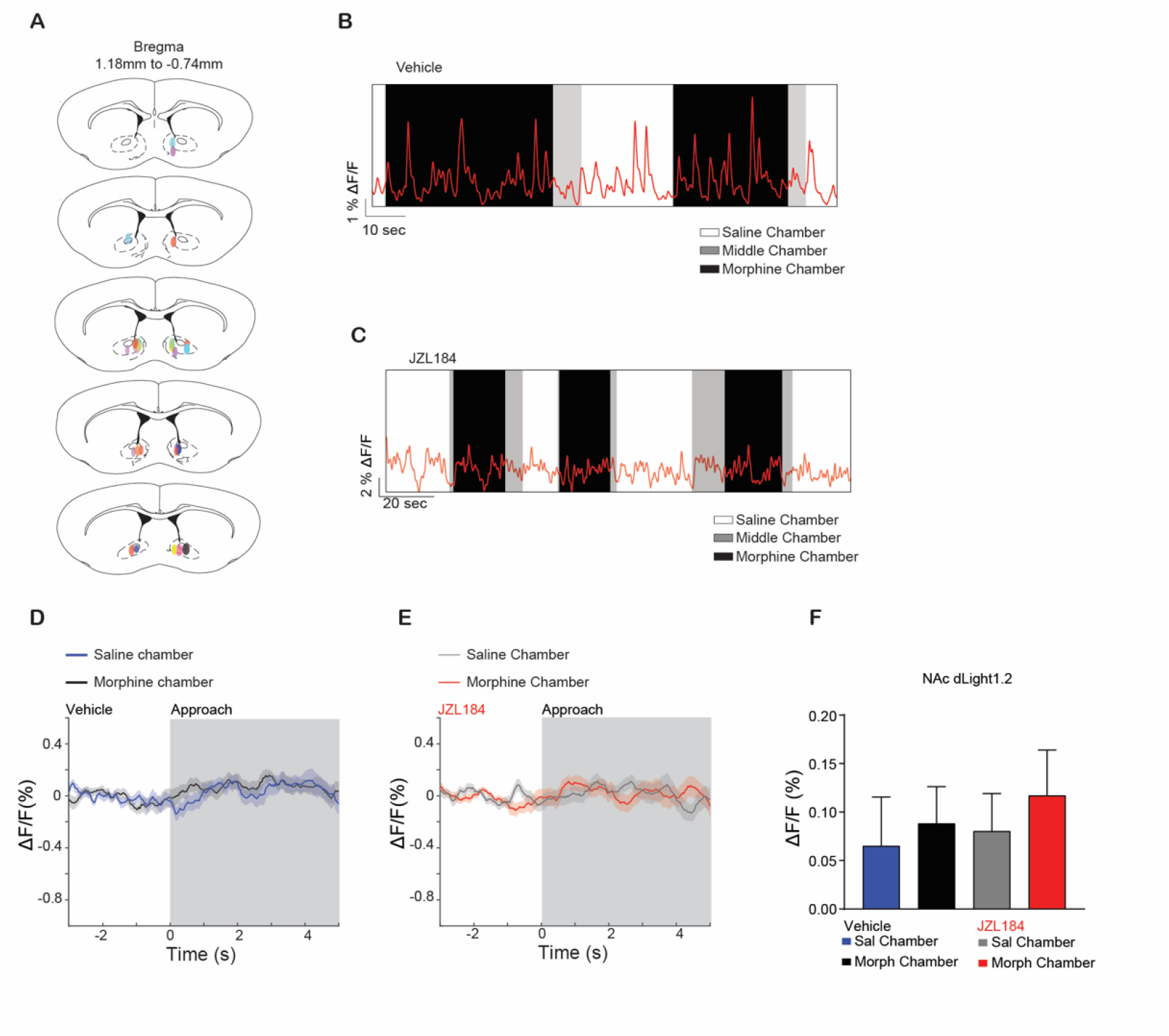
Brain placements and behavioral NAc dLight1.2 fiber photometry recordings. (**A**) Coronal sections of the fiber implant and dlight1.2 viral expression in the NAc. (**B**-**C)** Representative dlight1.2 traces collected during the CPP preference test day 6 in vehicle (**B**) or JZL184 (**C**) pretreated animals. (**D**) A representative trace of the mean ΔF/F (+/– SEM) photometry signal of NAc dLight1.2 fluorescence in vehicle pretreated mice time-locked to the exit from saline-paired chamber approaching the morphine-paired chamber (black) or exit from morphine chamber approaching the saline chamber (blue). (**E**) A representative trace of the mean ΔF/F (+/– SEM) NAc dLight1.2 fluorescence signal in JZL184 pretreated mice time-locked to the exit from saline-paired chamber approaching the morphine-paired chamber (red) or exit from morphine chamber approaching the saline chamber (grey). (**F**) No significant difference was observed in vehicle or JZL184 pretreatment groups (linear mixed effects model, Vehicle: p=0.7145, number of approaches = 588, n = 6; JZL184: p=0.5267 number of approaches = 523, n = 8).

